# Development of a multifunctional toolkit of intrabody-based biosensors recognizing the V5 peptide tag: highlighting applications with G protein-coupled receptors

**DOI:** 10.1101/2023.02.05.527200

**Authors:** Manel Zeghal, Kevin Matte, Angelica Venes, Shivani Patel, Geneviève Laroche, Sabina Sarvan, Monika Joshi, Jean-François Couture, Patrick M. Giguère

## Abstract

Protein-protein interactions (PPIs) form the underpinnings of any cellular signaling network. PPIs are highly dynamic processes and often, cell-based assays can be essential for their study as they closely mimic the biological intricacies of cellular environments. Since no sole platform can perform all needed experiments to gain a thoroughly comprehensive understanding into these processes, developing a versatile toolkit is much needed to address this longstanding gap. The use of small peptide tags, such as the V5-tag, has been extensively used in biological and biomedical research, including labeling the C-termini of one of the largest human genome-wide open-reading frame collections. However, these small peptide tags have been primarily used *in vitro* and lack the *in vivo* traceability and functionality of larger specialized tags. In this study, we combined structural studies and computer-aided maturation to generate an intracellular nanobody, interacting with the V5-tag. Suitable for assays commonly used to study protein-protein interactions, our nanobody has been applied herein to interrogate G protein-coupled receptor signalling. This novel serviceable intrabody is the cornerstone of a multipurpose intracellular nanobody-based biosensors toolkit, named iBodyV5, which will be available for the scientific community at large.

## INTRODUCTION

Conventional antibodies have proven to be robust workhorses in various cell biology research fields ^1^. Notwithstanding their indispensability, their large size and complex architecture preclude them from functioning within living cells ^2^. These limitations spurred advances in engineering novel recombinant antibody constructs, termed intracellular antibodies or intrabodies. Although their biophysical properties are unlike the conventional monoclonal antibodies, intrabodies retain the capacities for recognition and/or neutralization with similarly high specificities and affinities, with the added conferred ability to bind specific targets expressed within the same living cell that houses it, typically within the cytoplasm of eukaryotic cells ^3^. Despite intense efforts, minimal antigen-binding fragments such as single-chain variable fragments (scFvs) or F(ab)-type fragments remain highly unstable in a cellular environment and have limited activity in an intracellular milieu ^4-6^. The challenges for intrabodies to be expressed in their functional forms are partly due to their aggregation propensity and the reducing environment of the cytoplasm, which prevents intradomain disulfide bond formation ^7,8^.

Of greater promise are single-domain antibodies (sdAbs), also referred to as VHHs or nanobodies (Nbs), which are the smallest antibody derivatives (12-15 kDa). Although they were originally derived from members of the *Camelidae* and *Chondrichthyes* family, human-derived sdAbs have also been engineered from conventional human IgGs ^9^. Epitope-specific Nbs can be selected by various approaches, including display-based panning or synthetic libraries generated *in vitro* using predesigned scaffolds that undergo CDR codon randomization ^10^. Moreover, retrieved Nbs can be affinity-matured to further improve upon the original pharmacological features ^11^. Structurally, these non-canonical antibodies are more compact, consisting exclusively of shortened heavy chains with three interspersed complementarity-determining regions (CDRs) that form their antigen-binding domain. Devoid of the hydrophobic interface between heavy and light chains and stabilized by only a single internal disulfide bond, Nbs rival conventional IgGs given their low immunogenicity, enhanced solubility, and stability, all while retaining high binding affinities towards their targets, and thereby are more successfully expressed in the cytosol of mammalian cells ^12^. In terms of binding performance, Nbs have an enhanced ability to target sterically hindered and/or concave epitopes given their slightly convex-shaped paratope ^13^. As such, the physicochemical and pharmacological properties of Nbs offer a multitude of possibilities for live-cell applications, such as elucidating signalling pathways and protein-protein interactions in live cells, targeting proteins or protein complexes formerly inaccessible to biologics, fusing to other functional effectors such as proteases and fluorophores, visualizing and localizing proteins of interest using conventional and advanced microscopy ^14^. Nb targeting intracellular targets are plentiful, the majority of which target various cytosolic proteins ^15-28^.

### Study of protein-protein interactions

Different experimental techniques are commonly used to detect or isolate protein complexes, such as pull-down and immunoprecipitation. However, these approaches are disruptive to larger complexes, including membrane-spanning proteins. However, measurement and detection of protein-protein interactions is challenging and requires technology capable of measuring both high- and low-affinity interactions and transient and stable complexes. Despite the value of cell-based assays, the imbroglio of the protein interactome can complicate the execution of its study. Moreover, protein-protein interaction (PPI) assays, such as resonance energy transfer or binary complementation, require cloning of each component in all potential and permissive orientations, specifically at the C- or N-terminus of the protein, which could translate into over twelve different combinations to be tested. Finally, studying protein-protein interactions at a scaled-up level using cell-based assays has been limited thus far.

### Protease-dependent reporter assay (TANGO)

Protease-dependent reporter assays such as the Tobacco Etch protease-dependent assay (TANGO) are very sensitive and versatile assays that can measure the interaction between two proteins. Previous reports have exploited this assay to probe β-arrestin recruitment at select GPCRs or performing high-throughput screens of the entire class A GPCR-ome simultaneously ^29-31^. This assay relies on a TANGO-ized GPCR carrying a transcription factor at its C-terminus (tTA) preceded by a tobacco etch virus endopeptidase (TEV) cleavage site. Once the β-arrestin fused with the TEV protease (β-arrestin-TEV) is recruited to the receptor, it cleaves its recognition site and releases the tTA transcription factor, which translocates to the nuclease and activates its reporter gene under the control of the tTA-response element (TRE). Recently, a truncated version of the TEV (TEV219) has been demonstrated to be superior to its WT counterpart given its improved efficiency and that it does not self-inactivate; as such, TEV219 was selected to develop an updated GPCR-TANGO platform (submitted publication).

### Bioluminescence Resonance Energy Transfer (BRET)

Biophysical techniques involving resonance energy transfer enable monitoring of the formation of dynamic complexes in living cells ^32^, with their use being well-established and remaining a gold standard in GPCR signaling ^33^. The receptor is fused at the C-terminus with the *Renilla* Luciferase variant 8 (RLuc8)^34^ and the nanobody is fused at either N- or C-terminus with the GFP2 acceptor ^35,36^.

### Nanoluciferase Binary Technology (NanoBiT)

The NanoBiT assay is a structural complementation reporter system composed of a Large BiT (LgBiT; 18kDa) subunit and a small complimentary peptide named Small BiT (SmBiT; 11 amino acid peptide), which has been optimized to have low affinity for LgBiT. When the two proteins interact, the subunits assemble into an active enzyme which, in the presence of its substrate, furimazine, results in the production of a luminescent signal in living cells, enabling real-time kinetic measurements of protein interaction dynamics ^37^. The typical configuration consists of the receptor fused at the C-terminus with the SmBiT peptide, which with respect to GPCRs, likely minimizes the receptor’s native state or ligand-induced complexes.

Although these approaches have been designed to provide an understanding of different PPIs, all require the fusion of large, functionalized tags (TEV, GFP2, LgBiT) to the proteins of interest. Consequently, the small size of peptide tags (<20-mer) could be preferable for these purposes instead of using bulkier fluorophores and polypeptides whose addition could interfere with the native behaviour and functions of the protein of interest such as complex formation and regulation, especially interactions that are particularly proximity- or location-dependent ^38^. Besides avoiding cross-reactions within the native proteome, epitope tags obviate the need for custom antibody generation for every protein of interest and give rise to the possibility of multiplexing orthogonal peptide tags within the same experiment ^39,40^. Although epitope tags are versatile, Nbs recognizing these small peptide fragments are scarce, in part due to conformational difficulties of the cleft-like paratope of Nbs for recognizing linear peptides and the needed strength of Nb-peptide tag interaction. Based on existing literature, less than 10 peptide-targeted Nbs have been identified and few were used as intracellular nanobody (intrabody)^41,42^.

Therefore, the paucity of tag-targeted intrabodies is a bottleneck for intracellular interrogations, especially given the pre-existing arrayed or pooled tag-encoded cDNA and ORF libraries for overexpression studies. For example, while there are comprehensive human genome-wide and fully annotated V5-tagged libraries developed from the ORFeome project ^43^ and the *Drosophila* genome-wide V5-tagged library ^44^, these open resources are mainly limited to phenotypic screenings for living cellular assay or microscopy for *in vitro* assay on fixed tissues.

The V5 tag is a universal epitope that has been extensively used since its introduction roughly 30 years ago ^45^. In 1987, a monoclonal antibody binding the P and V protein of the simian virus 5 (SV5) was isolated and called SV5-Pk ^46^. The epitope recognized by the SV5-Pk was later identified as the residues 95-108 of the RNA polymerase alpha subunit and was subsequently termed the “V5-tag”, with the sequence of this peptide being GKPIPNPLLGLDST ^45-47^ (**Fig. 1a**). The strong affinity of this antibody for the 14-mer peptide tag was later utilized to generate a purification platform and is now commonly applied for immunodetection of V5-tagged proteins by Western Blot or immunohistochemistry. This small (1.42 kDa) tag contains an equal number of positively and negatively charged residues. Additionally, its prokaryotic origin circumvents possible background signals when expressed in mammalian and insect cell hosts and is also resistant to cleavage by their endogenous proteases ^42^.

**Fig 1:**
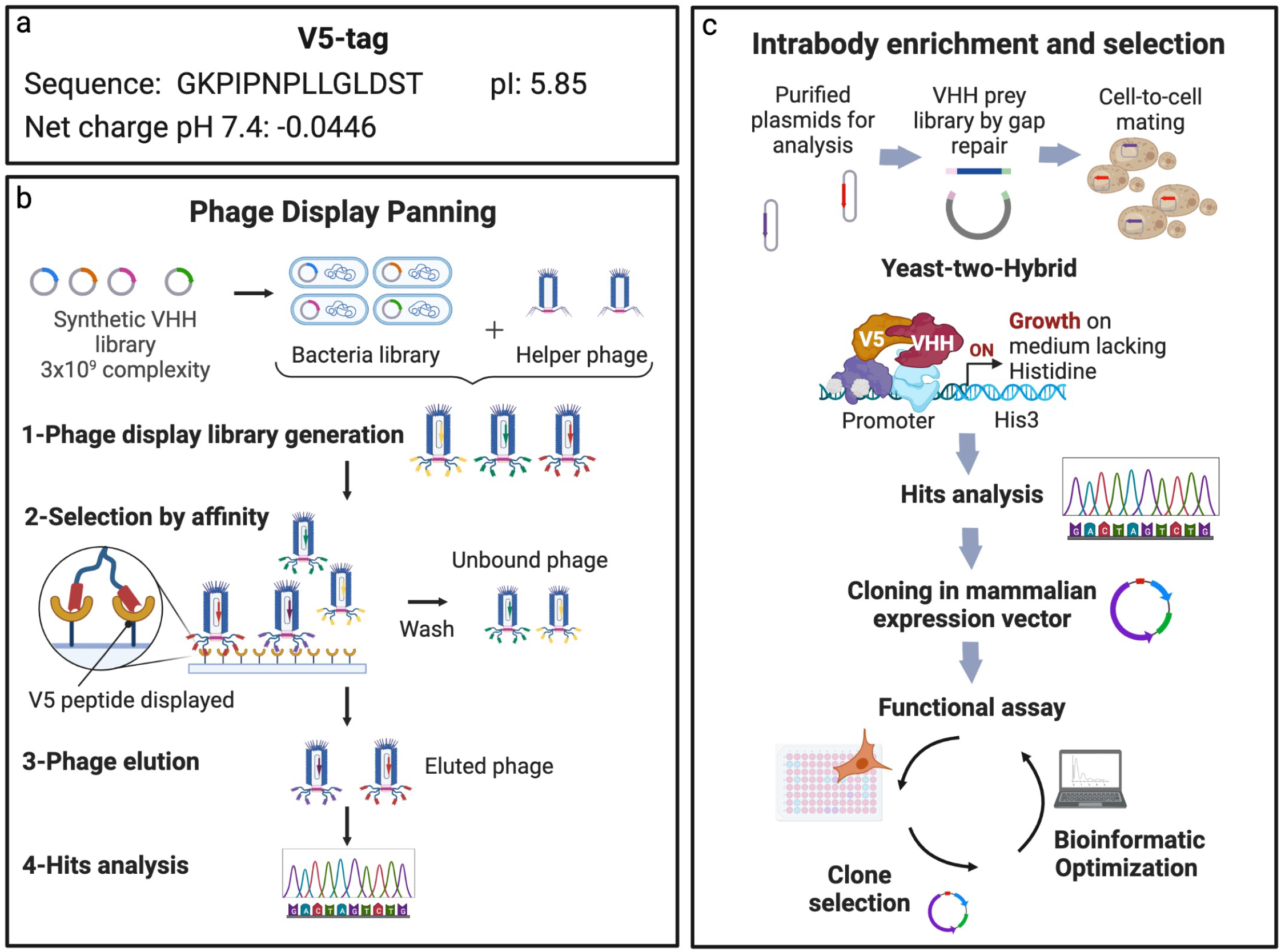
Overview of the selection of a synthetic nanobody interacting with the V5-tag. **a**, Sequence and biochemical properties of the V5-tag. **b**, Schematic of the phage-display planning for the enrichment of nanobodies interacting with the V5-peptide tag. **c**, Schematic of the yeast-two-hybrid screening (Y2H) for the enrichment of nanobodies interacting with the V5-peptide tag in an intracellular environment (intrabody).

Therefore, the development of intracellular Nb recognizing the small peptide V5-tag and its functionalization would be highly amenable to cellular studies of a broader and systematic nature (genome-wide), as well as more directed investigations of gene function and genetic rescues, and protein-protein interactions such as dynamic receptor-protein interfaces, with minimal interference. To this end, we introduce the NbV5, as a novel nanobody that interacts with the V5-tag, which serves as the foundation of our multipurpose toolkit comprising intracellular nanobody-based biosensors, dubbed iBodyV5.

## RESULTS

### Generation of a nanobody directed against the V5-tag

The first generation of selective nanobodies recognizing the V5-tag were identified and selected for using the Hybribody platform. Briefly, this approach combines synthetic VHHs phage-display screening and yeast two-hybrid (Y2H). For our application, CDRs were inserted into humanized scaffolds to generate the synthetic library NaLi-H1 ^48^. This combined strategy has the advantage of enriching stable and soluble intracellular-targeting nanobodies; the overall process is illustrated in **Fig.1b-c**. The peptide phage display was first performed against a purified HaloTag protein carrying a duplicate copy of the V5-tag. The VHHs selected after one round of phage display were cloned into yeast prey vector to perform yeast-two hybrid screening ^49,50^, finally obtaining 52 different VHHs with redundancies ranging from 1 to 37. The first 20 clones with the most redundancies were selected, and VHHs were cloned into pcDNA3.1^+^ fused at the C-terminus with the TEV219. Each clone was assayed using the TANGO-based assay using the mu-opioid receptor (μ-OR) and Angiotensin II receptor type 1 (AT_1_R) and the β-arrestin-2 carrying a N- or C-terminal V5-tag (V5-β-arrestin-2 and β-arrestin-2-V5, respectively). The overall schematic of the nanobody-based TANGO assay is depicted in **Fig.3a**. Each clone was tested independently using an increasing concentration of the μ-OR agonist DAMGO and AT_1_R agonist angiotensin II as previously described ^29-31^. From these preliminary findings, only one clone referred to herein as NbA1, showed a powerful response with β-arrestin-2-V5 but a feeble response toward V5-β-arrestin-2. Notably, codon optimization for human expression of the original nanobody resulted in a 3-fold increase in expression and was selected for further experiments in human cells (**Fig.3 b-c**).

**Fig. 2:**
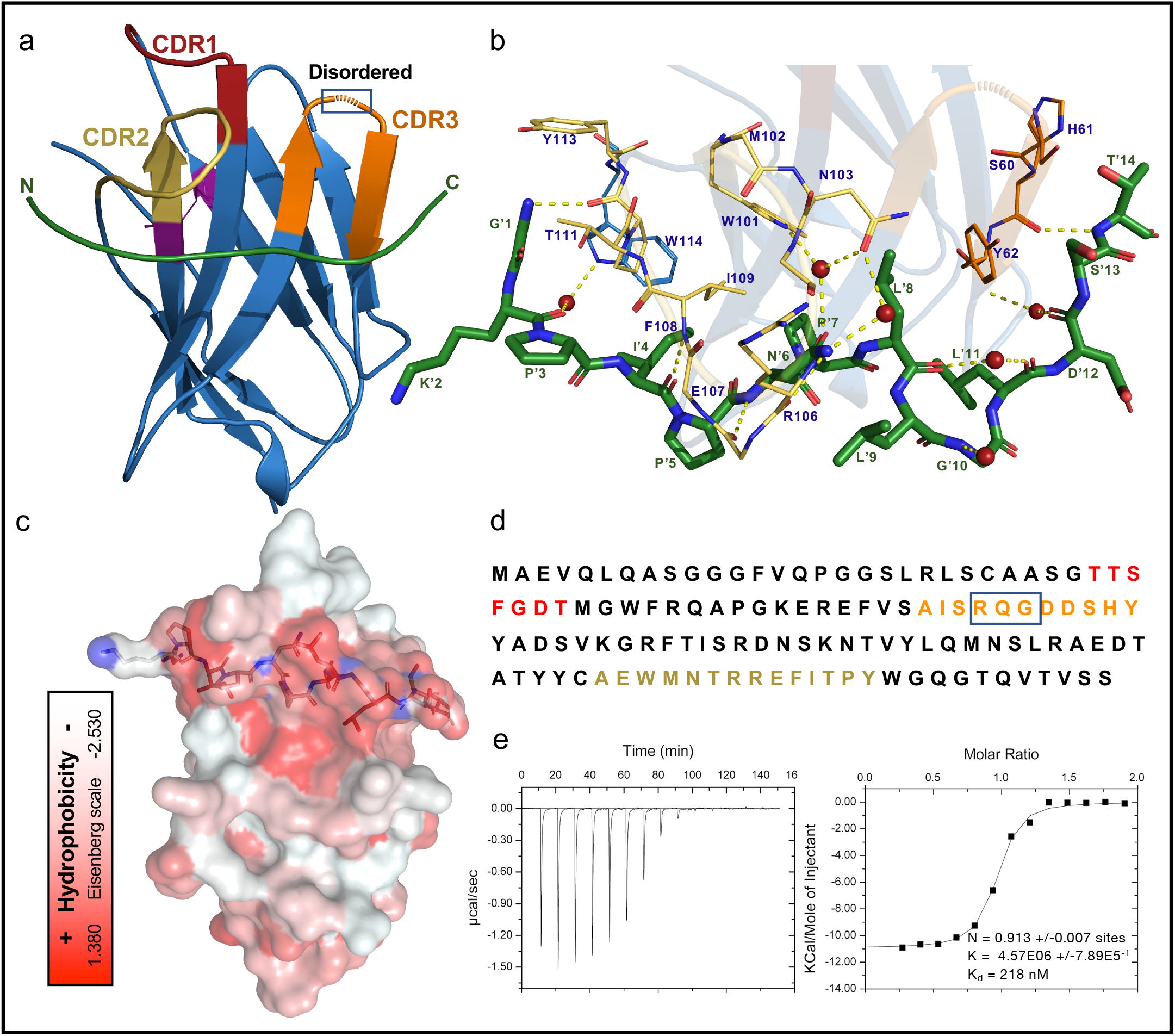
Structure of the NbA1 bound to the V5 peptide. **a**, Overview of the NbA1-V5 peptide complex shown as cartoon representation. NbA1 is colored in blue with CDRs 1-3 colored in red, orange, and yellow, respectively. Three residues from the CDR2 were not resolved in the refined structure. **b**, Close-up view of the polar interactions within the complex including water-bridged interaction. The V5-peptide (green) is shown as stick representation with the N-terminal on the left. Interacting residues from the paratope are labeled in blue while the peptide labels are green and marked with a prime symbol. H-bonds are shown as yellow-dashed lines and water molecules as red spheres. **c**, The NbA1-V5 peptide complex shown as surface representation and colored using the color_h script based on the Eisenberg hydrophobicity scale ^64^. **d**, Sequence of NbA1 with CDRs 1-3 colored in red, orange, and yellow, respectively. The square highlights the RQG tripeptide that is disordered in the CDR2 loop. **e**, Isothermal titration calorimetry curve of V5-peptide tag titrated into NbV5 in absence of DTT and corresponding ITC values. Titrations were performed in duplicate. S.D. represents the standard deviation between the two experiments.

**Fig. 3:**
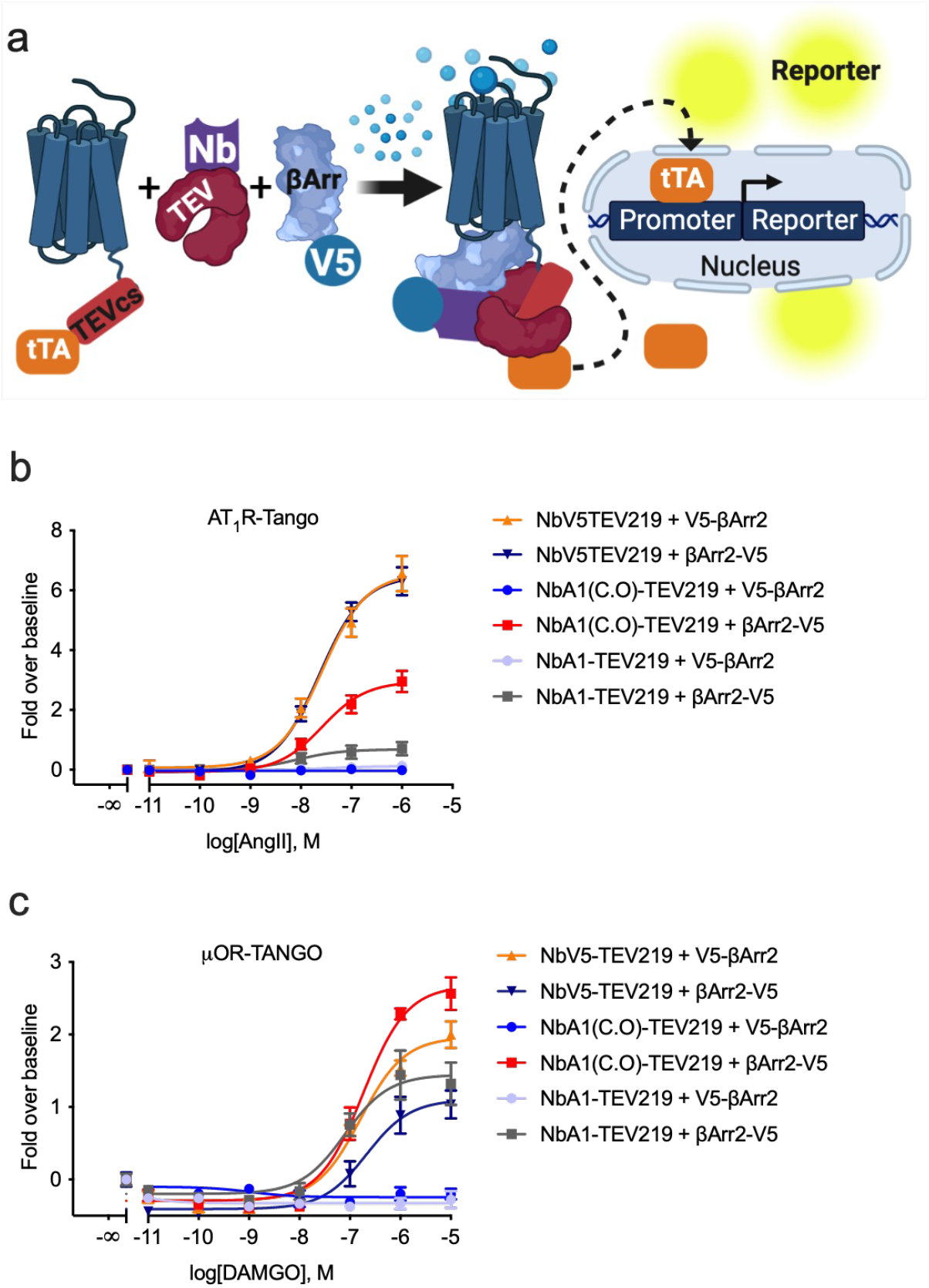
NbV5 as a versatile nanobody-based biosensor: application in protease-dependent cell-based assay (TANGO). **a**, Schematic of the protease-dependent cell-based assay (TANGO) used to assay the anti-V5 nanobodies. The original NbA1 clone selected from the Y2H screening was fused with the TEV219 protease and cloned into a eukaryotic expression vector. The initial assessment revealed that the nanobody was not well expressed in HEK293 and codon optimization (NbA1(C.O)) increased its expression and consequently its functional activity. The AT_1_R-TANGO (**b**) and μ-OR-TANGO (**c**) were co-transfected with β-arrestin1 or β-arrestin2 carrying a C- or N-terminal V5-peptide tag along with either the NbA1-TEV219, NbA1(C.O)-TEV219 or NbV5-TEV219 fusion protein. Dose-response agonist treatments demonstrate the difference between the NbA1 and NbV5 nanobodies. The NbV5 recognizes the N- or C-terminally V5-tagged β-arrestin2 with similar logistic parameters (potency and efficacy) while the original NbA1 clone only recognizes the C-terminally tagged β-arrestin2 (β-arrestin2-V5). Dose-response curves were built using XY analysis for non-linear regression curve and the 3-parameters dose-response stimulation function. All error bars represent SD (n = 3 technical replicates).

### Crystal structure of the NbA1

Given the suboptimal activity of NbA1 toward the N-terminally tagged β-arrestin-2, we thus opted for a computer-aided affinity maturation approach; to do so, we first solved the V5-bound structure of NbA1 at a resolution of 2Å (**Table 1**). NbA1 shapes as a conventional V-shape IgG fold connected by a disulfide bond while the V5-tag does not have any secondary structures ^51^ (**Fig.2a**). The paratope in which the V5 peptide interacts is primarily mediated by the CDR2 and CDR3 regions, which is common for nanobodies ^52^ (**Fig.2a**). The V5 peptide makes several polar and hydrophobic contacts with NbA1 and is orientated at 90 degrees to the central axis of the nanobody. Out of the 14 residues of the tag, 11 are directly interacting with NbA1 (**Fig.2b**). the lateral chains of two residues, Lys 2 and Ile 4, point outward but their backbones make water-mediated hydrogen bonds with NbA1. As shown in **Fig. 2c**, the interaction is mediated in large by hydrophobic residues, suggesting a strong contribution of entropic forces by the Nb:V5 interfaces ^53^. Nanobodies are known to have greater paratope diversity compared to classical Ab VH domains. This is highlighted by a recognition mode that is very different compared to the ALFA-tag, which involves contacts with all three CDRs and is oriented parallel to the central axis (**Ext. Fig.1**) of the nanobody. Finally, three residues (Gln^57^, Gly^58^, Asp^59^) within the CDR2 loop are disordered and were not resolved (**Fig.2a and Ext. Fig.1**). Further characterization of NbA1 was performed using isothermal titration calorimetry (ITC), which estimates the binding affinity of NbA1 to be 500 nM (data not shown).

**Table 1:**
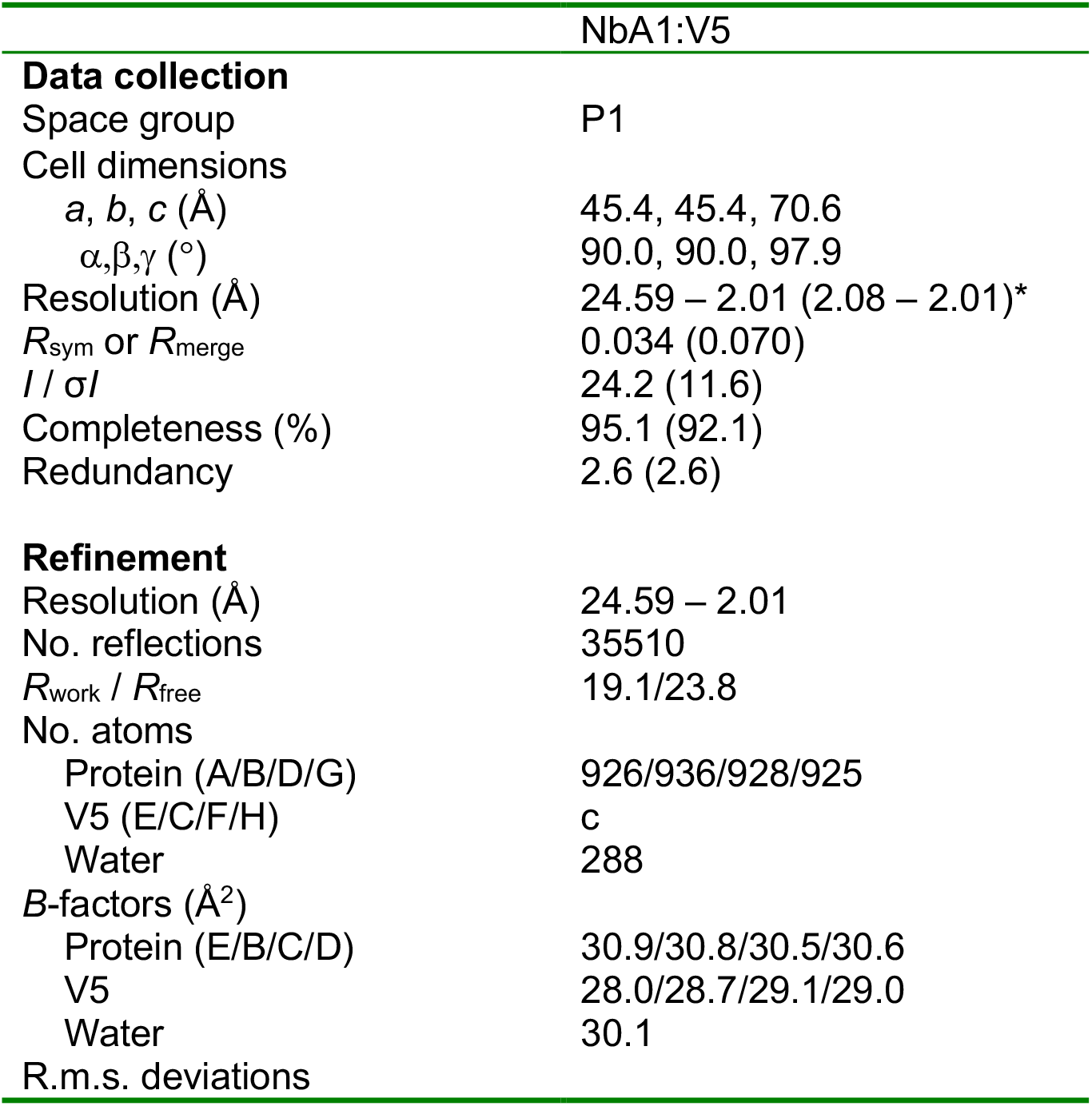
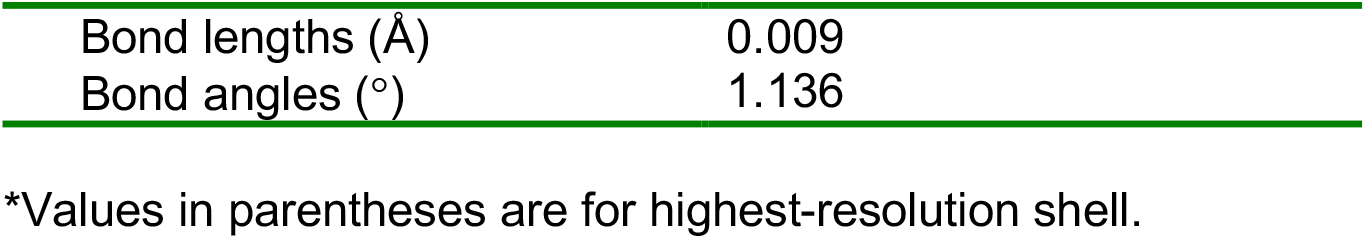
Data collection and refinement statistics for NbA1 in complex with the V5 peptide.

### Maturation of the NbA1

A major goal of our approach is to facilitate the study of protein-protein interactions using a versatile nanobody-based sensor. While NbA1 is functional, it showed a pronounced preference for the C-terminally tagged β-arrestin-2. This partiality may be explained by the lack of interactions with the last two residues of V5. To improve the overall affinity, especially for the N-terminus of V5, *in silico* affinity maturation was performed using the Rosetta modelling software. The relaxed model generated ten designed models, and the corresponding scores and binding energy metrics were plotted. Ten mutants with degenerated mutations were synthesized and tested in TANGO assay. Based on functional evaluation, it was deduced that two mutations (ΔD59 and S60K) resulted in better activity compared to NbA1, especially for the N-terminal V5-tagged arrestin. Consequently, an optimized nanobody was generated to incorporate only those two mutations, called hereafter NbV5. Notwithstanding the exact mode of interaction, this maturated NbV5 showed a major improvement in its overall behaviour in our functional TANGO assay (**Fig.3b-c**) and was selected moving forward to develop the toolkit. Moreover, based on ITC, the binding affinity of NbV5 was improved to an estimated 218 nM (**Fig. 2**).

The interaction between two proteins can be generalized as the “bait” and “prey.” In our platform, the bait is a membrane receptor belonging to the G protein-coupled receptor (GPCR) family. In the specific case of GPCRs, their membrane configuration only allows for C-terminal fusion. The prey protein can be any V5-tagged protein in which the tag is placed at the C- or N-termini as well as within the protein sequence. As a proof of concept, we used the well-established GPCR interactors β-arrestin-1 and β-arrestin-2 as the prey. In this tripartite system, the iBodyV5 will bridge the V5-tagged prey protein with the bait receptor carrying the other functional moiety, reconstituting a functional cell-based assay with minimal interference onto the prey protein’s function and the native complex formation.

The initial characterization of our platform with GPCRs allowed for regulation of the interaction using an agonist and thus, a better approach to measuring the efficiency of our biosensor approach. Subsequent demonstrations were conducted using two divergent GPCRs to characterize the iBodyV5 biosensors, specifically the angiotensin II receptor (AT_1_R) and the mu-opioid receptor (μ-OR), two widely used receptors in our lab.

### iBodyV5@TANGO biosensors

The first functional assay adapted with our tripartite system was the protease-dependent reporter assay. As described above, TANGO was used as our primary assay to assess our nanobody during the optimization because of its high sensitivity and permissibility. Further characterization at the μ-OR and AT1R showed a dose-response curve with both β-arrestin-1 and -2, as well as in both configurations with respect to the position of the V5-tag (**Fig.3**). It is important to mention that the EC50 is comparable to our updated TANGO assay with the TEV219 directly fused at the C-terminus of the β-arrestins (submitted publication). These results support that our iBodyV5 biosensors are well suited for use with the TANGO-based assay.

### iBodyV5@BRET biosensors

Another common functional assay that was adapted for our NbV5-based biosensor toolkit was BRET (**Fig.4a**). As for the TANGO, we compared both β-arrestin-1 and β-arrestin-2 fused at the N- or C-terminus with the V5-tag; the ALFA-tag was also included as a reference because the corresponding NbALFA intrabody is already characterized and was shown to work well with all placements (N-term, C-term, and internal) of the tag. As shown in **Fig.4b-d**, all configurations produced a robust signal, but C-terminally tagged NbV5 (NbV5-GFP2) gave a better BRET^2^ ratio, while the NbALFA is less affected. As shown in **Ext. Fig. 1**, the binding mode of NbALFA is at a 90-degree angle compared to NbV5, which could explain this observation. The placement of the tag on the prey protein does not seem to have a significant impact regarding the tested receptor-β-arrestin coupling, and for AT_1_R specifically, both β-arrestin isoforms are well recruited with a slight preference for β-arrestin-2 in this assay. Our iBodyV5 biosensors are thus favorable for application with BRET assay.

**Fig. 4:**
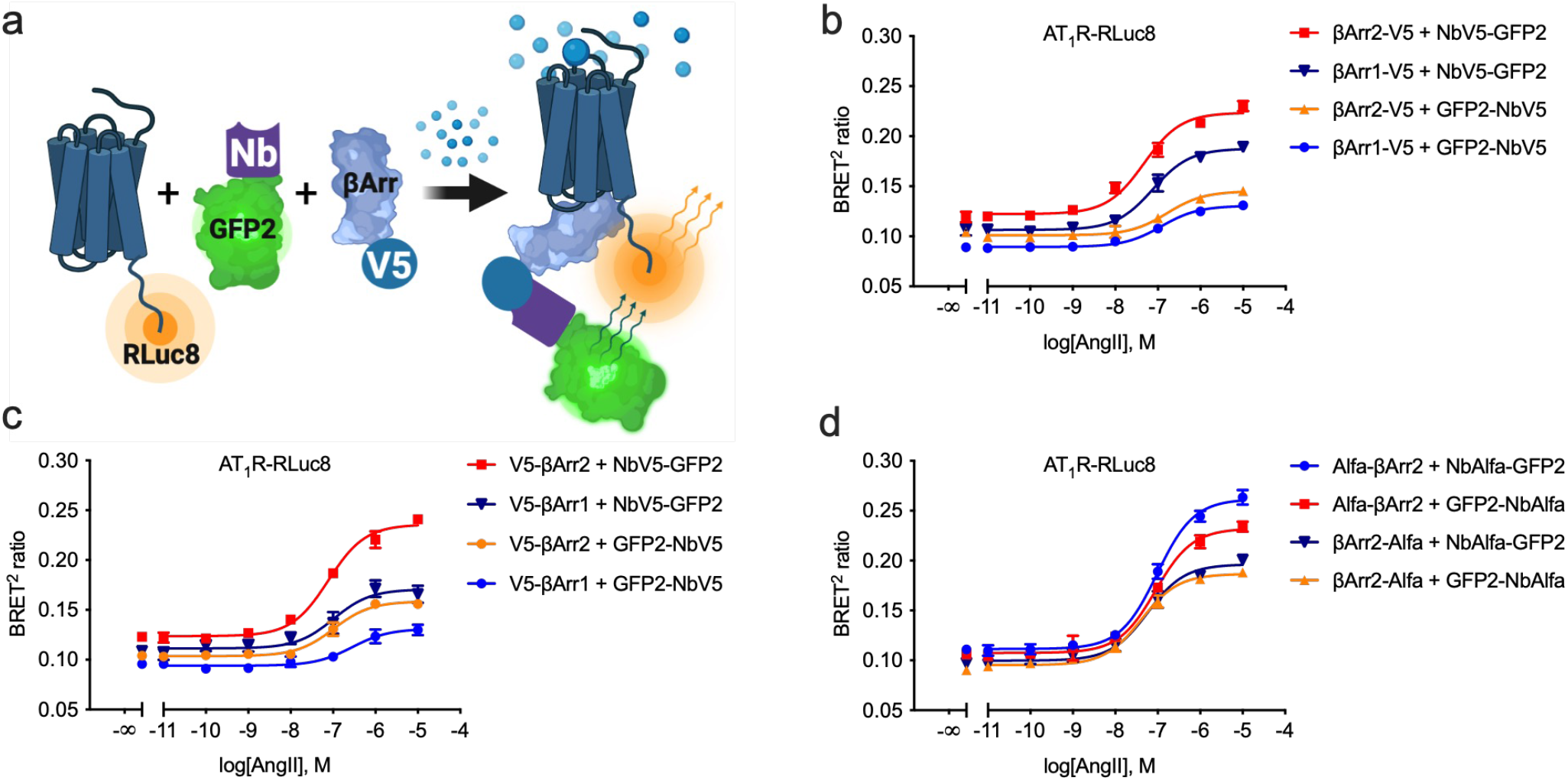
NbV5 as a versatile nanobody-based biosensor: application in *B*ioluminescence *R*esonance *E*nergy *T*ransfer (BRET^2^). **a**, Schematic of the BRET^2^ cell-based assay used to evaluate the NbV5. The recruitment of the β-arrestin-NbV5-GFP2 complex to the receptor fused to RLuc8 allows the energy transfer between the *Renilla reniformis* Luciferase mutant (RLuc8) and the green fluorescence protein mutant GFP2 in the presence of the RLuc8 substrate Coelenterazine 400a. **b-d**, NbV5-based detection of β-arrestin1 and β-arrestin2 recruitment at the AT_1_R-RLuc8. N- and C-terminally V5-tagged β-arrestins were tested as well as both N- and C-terminally GFP2-tagged NbV5. Dose-response curve treatment with angiotensin II reveals equivalent recruitment of β-arrestin1 and 2 at the AT_1_R. No major differences were observed between the N- and C-terminally V5-tagged β-arrestin, whereas the C-terminally GFP2-tagged NbV5 (NbV5-GFP2) showed a better response. **d**, Similar results were obtained with the ultra-potent nanobody that recognizes the synthetic ALFA-tag (NbALFA). Dose-response curves were built using XY analysis for non-linear regression curve and the 3-parameters dose-response stimulation function. All error bars represent SD (n = 3 technical replicates).

### iBodyV5@NanoBiT biosensors

Finally, we also introduced our nanobody biosensors for the nanoluciferase binary complementation technique. Varying configurations for both β-arrestins (V5-β-arrestin and β-arrestin-V5) and nanobody fusions (Nbs-LgBiT and LgBiT-Nbs) were compared (**Fig.5**). As opposed to the BRET^2^ experiment (**Fig.4)**, the LgBiT-NbV5 produced a better response compared to NbV5-LgBiT, which highlights the importance of testing both orientations for optimal response and sensitivity. One observation using NanoBiT is that, compared to the BRET^2^ assay, β-arrestin-1 is strongly favoured over β-arrestin-2 at the AT_1_ receptor (**Fig.5b-c**). Similar results were obtained with the ALFA-tag system, indicating the tag system does not affect the results (**Fig. 5d**). For the μ-OR, we opted to co-transfect the GPCR kinase 2 (GRK2) which is known to increase β-arrestins recruitment for this receptor (see **Fig.6e-f**). As seen in **Fig. 5e and f**, the β-arrestin-1 is favored compared to β-arrestin-2 for μ-OR and the N-terminally V5-tag β-arrestins and N-terminal LgBiT-NbV5 are the optimal orientations for this receptor.

**Fig. 5:**
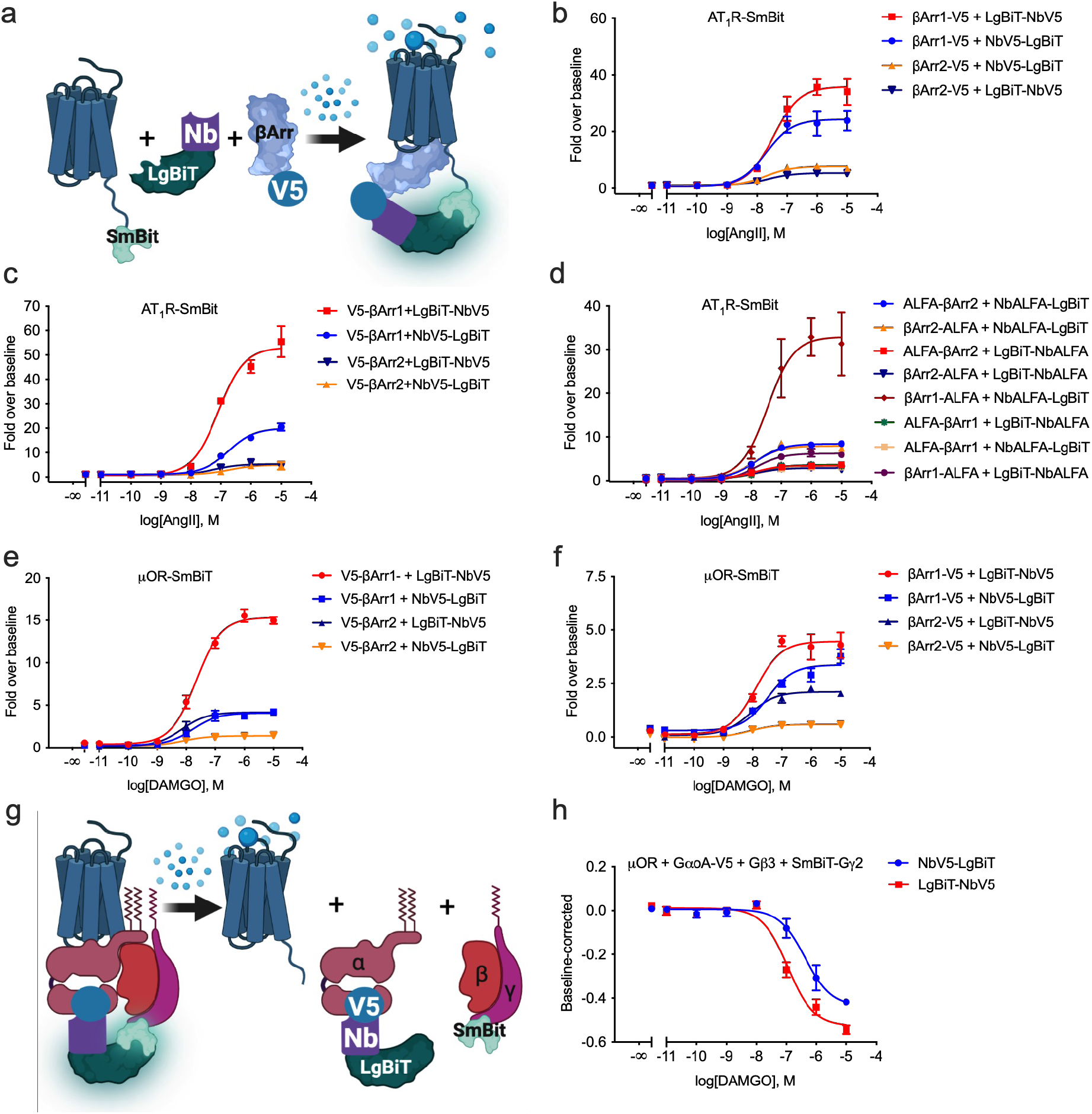
NbV5 as a versatile nanobody-based biosensor: application in *Nano*luciferase *Bi*nary *T*echnology (NanoBiT). **a**, Schematic of the NanoBiT cell-based assay used to evaluate the NbV5. The recruitment of the β-arrestin-NbV5-LgBiT complex to the receptor fused to the SmBiT allows the binary complementation of the active nanoluciferase enzyme, and the generation of a bright luminescent signal in the presence of its substrate furimazine. NbV5-based detection of β-arrestin1 and β-arrestin2 recruitment at the AT_1_R-SmBiT (**b**,**c**,**d**) and μ-OR-SmBiT (**e**,**f**) was assayed. N- and C-terminally V5-tagged β-arrestin was tested as well as both N- and C-terminally LgBiT-tagged NbV5. Dose-response curve treatment with angiotensin II shows the preference of AT_1_R for β-arrestin1 in this assay, which is contradictory with findings obtained with BRET^2^ (see discussion). In this assay/configuration, N-terminal LgBiT-tagged NbV5 (LgBiT-NbV5) shows a better response. Similar results were obtained with the ultra-potent nanobody that recognizes the synthetic ALFA-tag (NbALFA) (**d**) but in that configuration, the NbALFA-LgBiT produced a stronger signal. The NbALFA system produced a maximum ∼8-fold increase with β-arrestin2 which is similar to the NbV5 system. **g**, Schematic of the NanoBiT cell-based assay used to assess the internal V5-tag. The functional recognition of an internal localized V5-tag was also assayed by NanoBiT by incorporating the V5-tag at position 92 of the GαoA and measuring the dissociation from the Gβ3 and SmBiT-Gγ2 dimer upon μ-OR activation with DAMGO. Dose-response curves were built using XY analysis for non-linear regression curve and the 3-parameters dose-response stimulation function. All error bars represent SD (n = 3 technical replicates).

**Fig. 6:**
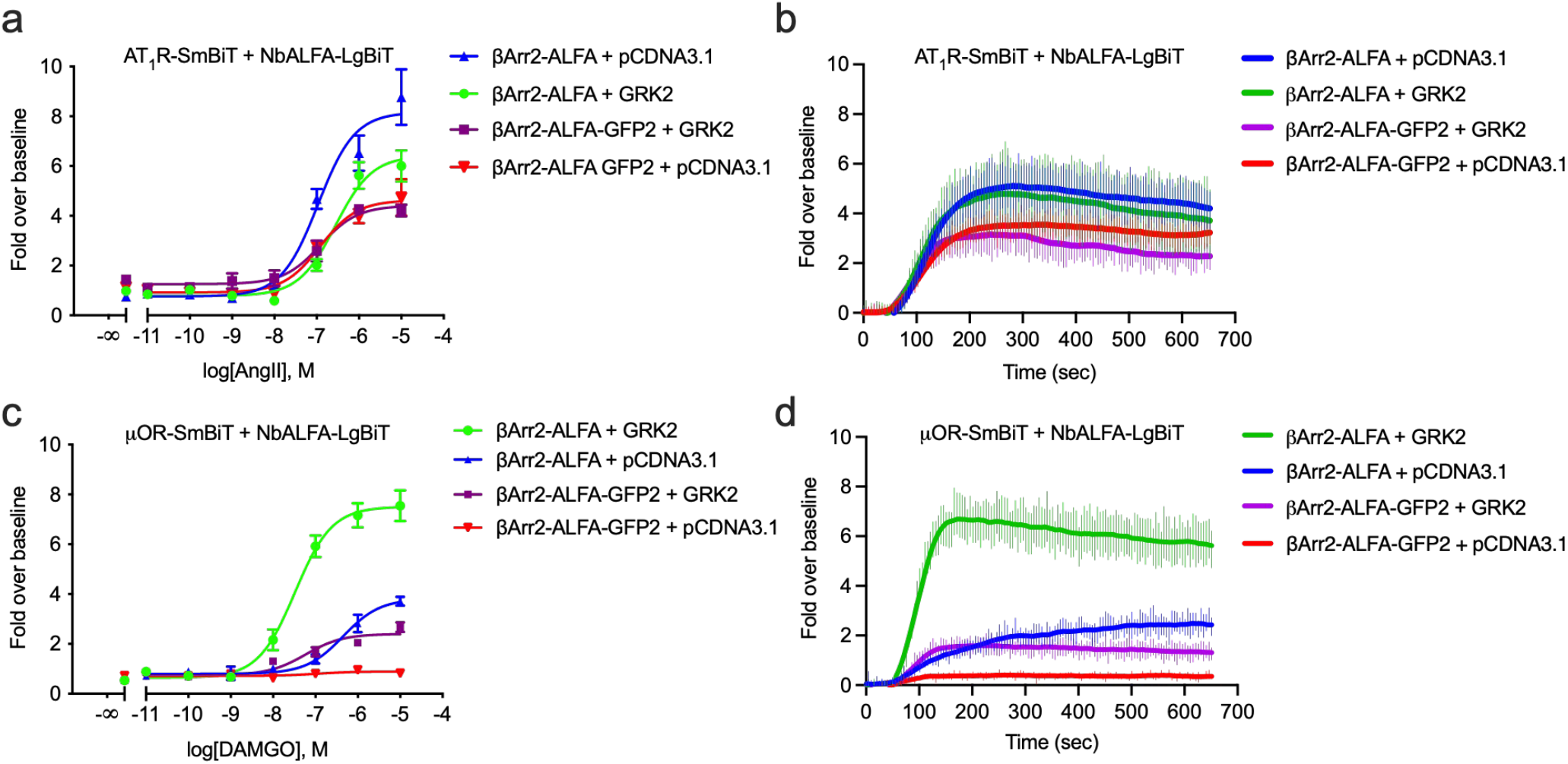
Effect of tag size on β-arrestin recruitment in BRET^2^ and NanoBiT assays. **a**, To assess the effect of a larger tag on the activity of β-arrestin2 recruitment at the AT_1_R, β-arrestin2-ALFA was compared against β-arrestin2-ALFA-GFP2. **b**, NbALFA-based detection of β-arrestin2 recruitment at the AT_1_R-RLuc8 was assayed in BRET^2^, with no difference being observed in this configuration. NbALFA-based detection of β-arrestin2 recruitment was then assayed in NanoBiT at the AT_1_ (**c**,**d**) and μ-opioid (**e**,**f**) receptors in the presence and absence of GRK2. The addition of GFP2 tag decreases β-arrestin2 at AT_1_R but also abolishes the cooperative effect of GRK2. A similar observation was made for the μ-OR but with more amplified phenotype. GRK2 has a strong cooperative effect on β-arrestin2 recruitment at the μ-OR. GFP2 almost completely abolishes β-arrestin2 recruitment at the μ-OR in the absence of GRK2 and strongly reduces the efficacy and potency in the presence of GRK2. Live kinetic trace at 1 μM agonist are shown in (**c**) for the AT_1_R and (**e**) for the μ-OR. GRK2 expression increases the rate of β-arrestin2 recruitment at μ-OR while having modest or no effect at AT_1_R. Dose-response curves were built using XY analysis for non-linear regression curve and the 3-parameters dose-response stimulation function. All error bars represent SD (n = 3 technical replicates).

Furthermore, the ability of NbV5 to recognize an internally positioned V5-tag was evaluated, specifically using the inhibitory G protein GαoA with the V5-tag inserted at position 92. It is well established that inhibitory G proteins tolerate insertion at this position, commonly used to insert a fluorescent protein or RLuc8 for BRET experiments ^54^. **Fig. 5g** illustrates the approach used, wherein the SmBiT was fused at the N-terminus of the G*γ*2 subunit and was co-expressed with Gβ3, assembling to make the obligatory Gβ*γ* dimer which interacts with the inactive GDP-bound GαoA. Once activated by the Gαi/o-coupled μ-OR, the activation of the G protein can be detected through the dissociation of the heterotrimer. As shown in **Fig.5h**, μ-OR activation by its agonist DAMGO induces a dissociation of the trimer that can be tracked using NbV5 fused with the LgBiT at N- or C-terminus. Thus, our Nb-based sensor allows the measurement of G protein activation via the insertion of the V5 sequence, which minimizes the possibility of affecting overall G protein regulation. Based on prior experiments performed in our laboratory, we have determined that when placed internally, NanoBiT complementary subunits are not well-tolerated, which is yet another advantage of using our V5-tagged biosensors.

All conventional assays use larger functionalized tags, such as GFP2, RLuc or TEV219 to name a few, fused to the prey proteins. Therefore, we aimed to measure the effect of the fusion of a larger tag to β-arrestin using the NbALFA:ALFA-tag system, with the assumption being that the rigidity of the ALFA-tag plus the inclusion of a proline before and after the tag would reduce the environmental impact.

The NanoBiT results (**Fig.5**) diverged from those obtained in BRET^2^ (**Fig.4**) in varying ways. Firstly, a preferential recruitment of β-arrestin1 compared to β-arrestin2 was observed, as well as a strong positional effect depending on the tag location. For this reason, we assessed the inclusion of a larger fusion tag using β-arrestin2-ALFA and β-arrestin2-ALFA-GFP2 at both AT_1_R and μ-OR, with the aim to see whether the GFP2 addition to the β-arrestin has any impact on its recruitment at the receptor. Hence, to detect reasonable recruitment at the μ-OR, the G protein-coupled receptor kinase 2 (GRK2) is often co-expressed, especially since its expression is absent at the endogenous level in HEK293 cells ^55^. As seen in **Fig.6c**, we observed a significant increase in β-arrestin recruitment at the μ-OR in the presence of GRK2. Even more striking is that the GFP2 tag at the C-terminus of the β-arrestin has a strong deleterious effect compared to that observed with AT_1_R in **Fig. 6b**. Interestingly, the presence or absence of GFP2 does not affect the EC50 (5.81E-06 and 3.3E-06 respectively), so the effect is presumably at the ‘‘quality’’ of the interaction owing to low-efficacy recruitment. We can therefore conclude that the bulkiness of the tag has an unpredictable effect, which depends on the bait:prey pair used, and should be avoided as much as possible. As such, the use of a tag system such as NbV5:V5 should minimize this potential effect.

One of the main advantages of the NanoBiT assay is the ease of performing a live experiment. While it can be used to perform endpoint assays on a microplate reader as in **Fig.6a and c**, with an appropriate real time kinetic reader, a live experiment can reveal additional information. This is especially true with GPCRs, as we assume that all interactions occur under equilibrium conditions, but it is well established, at least as with the case of heterotrimeric G proteins, that the interaction is short-lived ^56^. Thus, to study GPCR signaling in non-equilibrium conditions, kinetic experiments and appropriate models should be favorized ^57^. As such, we demonstrate in **Fig. 6b and d** a live kinetics experiment involving the recruitment of β-arrestin-ALFA versus β-arrestin-ALFA-GFP2 at the μ-OR and AT1R in the presence and absence of GRK2 at a single concentration of selective agonist. As previously published, the expression of GRK2 has the opposite effect at the AT_1_R compared to μ-OR^58^, specifically increasing the efficacy and potency at μ-OR but decreasing these parameters at AT_1_R. While the latter is well-represented in the endpoint experiment, the live experiment revealed that for the μ-OR, the expression of GRK2 increases the rate of β-arrestin recruitment dramatically and reaches a steady state that is relatively stable over time.

### Nanobodies as fluorescence trackers for microscopy

We next sought to determine whether the NbV5 fused with a constitutive fluorescent protein like eGFP could be used as a genetically-encoded tracker. Towards this, *γ*-actin carrying a C-terminal V5 tag (*γ*-actin-V5) was co-expressed with the NbV5-eGFP fusion protein. As shown in **Fig. 7**, the NbV5-eGFP diffuses within the cell in the absence of *γ*-actin-V5 but associates with *γ*-actin-V5 in its presence. Importantly, different actin-related structures like stress fibers and lamellipodia can be observed, indicating that the NbV5-eGFP could efficiently interact with a dynamic protein such as *γ*-actin without interfering with its natural localization. Therefore, iBodyV5 could be used to track protein remodeling or trafficking in a live-cell-based experiment.

**Fig.7:**
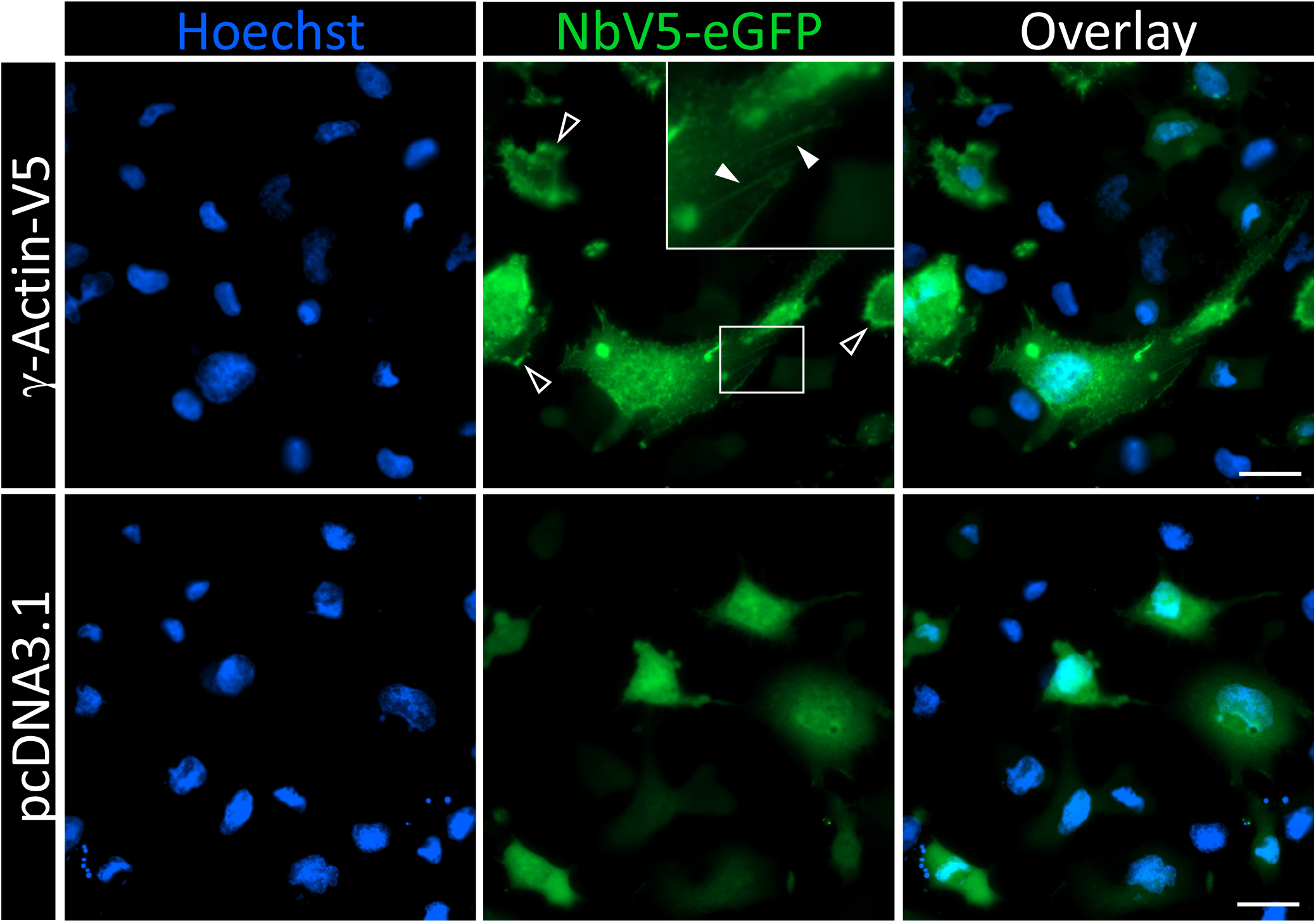
NbV5-based detection of V5-tagged proteins by immunofluorescence. Microscopy images of HT1080 cells expressing NbV5-eGFP and γ-Actin-V5 or pcDNA3.1+ alone as a negative control. In the absence of γ-Actin-V5, NbV5-eGFP is diffuse throughout the cell while in the presence of γ-Actin-V5, NbV5-eGFP is enriched in actin-rich protrusions and structures. For example, NbV5-eGFP is recruited to lamellipodia (open arrowheads) and stress fibers (closed arrowheads), indicating the complex does not alter the function of γ-Actin. Images are representative of 25 cells from three independent experiments. Scale bars are 100 μm.

## DISCUSSION

In this study, we are reporting the development and characterization of a V5-tag nanobody and a set of nanobody-based intracellular biosensors with multipurpose functionalities. The premise of our approach was to avoid the fusion of a large, functionalized tag to a protein and, more importantly, to harness an open-source genome-wide library of C-terminally V5-tagged ORFs. The V5-epitope is a well-established peptide tag that has been exploited for 30 years. The 14-mer peptide has no net charge at physiological pH and although it does not have any predicted secondary structures, it has quite an ordered core due to the presence of three proline residues. This intrinsic rigidity is believed to reduce the multidimensionality of the conformational landscape, thus minimizing the impact of environmentally induced secondary structures, as well as unpredictable behavior in solution. Many peptide tags have been developed over time, all with their respective advantages and disadvantages, yet few tag systems have corresponding functional intrabodies that are fully characterized as done in this study; the noteworthy exception is the NbALFA:ALFA-tag, the only well-characterized tag system on the market, although suffice to say that it is an artificial peptide tag. Although the ALFA works well, it is not an open-source resource which could limit further exploitation; moreover, the rigid helical structure of the ALFA tag could impact local secondary structures within cells. The availability of two different tag systems could also be capitalized on through simultaneous two-tag implementations.

Existing biotechnologies, used for the characterization and/or validation of potential drug-targets and the elucidation of physiological and pathophysiological cellular processes, require information often extracted from the abundance, localization, and dynamic interactions of cellular components^59^. As such, the development of intracellular traceable proteins for endpoint studies or real-time kinetics diagnostics is necessary, more so considering increasing efforts to obtain a more comprehensive understanding of these phenomena and thus expanding the feasibility of said intracellular tracers to a genome-wide or even to an organism (-omics-) scale.

Most nanobodies available to date were developed by animal immunization, but the development of synthetic *in vitro* platforms as used herein is of great promise ^48,60^. The Hybribody approach combines peptide phage-display and intracellular yeast-two-hybrid for further enrichment of potential intrabodies. Therefore, the selection of Nbs wherein the paratope folding does not rely on disulfide bonds is crucial for discovering potential intrabodies.

From this screening, potential nanobodies were isolated, with clone NbA1 functioning as the superior intrabody. Initial experiments revealed a very low level of expression of all nanobodies, which was circumvented by codon optimization for a human expression system. Further characterization in the TANGO cell-based assay showed that the NbA1 was not optimal for recognizing the V5-tag placed on the N-terminus of β-arrestin2. Thus, the NbA1 was purified from the periplasm of SHUFFLE bacteria, which are best suited for purification of proteins containing disulfide bonds. The purified NbA1 was co-crystallized with the V5-tag and the generated structure was used for computer-aided maturation to generate the final optimized NbV5 with improved *in vitro* functionality. The maturation was restricted to the three CDRs, with no mutants predicted to have major binding energy changes. Based on the mutants tested in TANGO, it was determined that two mutations improved the recognition of the V5-tag. The main advancement of the NbV5 compared to NbA1 is its improved interaction with the N-terminally positioned V5-tag (**Fig.3**). The two mutations were found to be juxtaposed wherein the Asp^59^ is removed and the Ser^60^ is mutated to a Lysine residue (ΔD^59^, S^60^K) (**Ext Fig.1**). As these two mutations border a disordered loop in the CDR2, the deletion of one residue may stabilize this region while Lys^60^ makes direct contacts with the CDR2 loop to further stabilize this region (**Ext Fig.1b, c**). The *in vitro* affinity was estimated at approximatively 218 nM, as measured using an isothermal titration calorimetry curve of V5-peptide tag titrated into NbV5 in the absence of DTT (**Fig.2e**). In the presence of DTT, which reduces disulfide bonds, the affinity was reduced to 574 nM (**Ext. Fig.2**). Given the reductive environment of the cytoplasm, the intracellular affinity is presumably between 200-600 nM.

Beyond the exact mode of interaction, we then moved forward to generate a comprehensive toolkit for versatile application in the most widely used cellular assays, namely TANGO, BRET and NanoBiT assays.

As shown herein, our nanobiosensors probed the recruitment of both β-arrestin-1 and β-arrestin-2 at two well characterized GPCRs, the μ-OR and the AT_1_R. The μ-OR is the receptor of the most prescribed analgesics such as morphine and fentanyl, while the AT1R is an important receptor for controlling vasocontraction and AT_1_R antagonists (ARBs) are prescribed for the treatment of hypertension, congestive heart failure and diabetic nephropathy. These two receptors are known to recruit both β-arrestins but there is some conflicting data regarding isoform selectivity and more importantly, the intrinsic drug efficacy. The measurement of the direct interaction between the receptor and its effector is a key aspect for the accurate determination of intrinsic drug efficacy and as such, the use of smaller tags should mitigate some distortions caused by larger functional tags.

The first assay successfully converted for inclusion in the NbV5-based toolkit is the TEV-dependent reporter method. This robust reporter assay was chosen for its permissibility and sensitivity. Despite these advantages, a potential limitation of TEV-dependant reporter methods is the kinetics of catalysis. In 2020, Sanchez et al. reported the S153N TEV219 mutant with an increased catalytic rate ^61^. We thus tested this new mutant and observed a 3 to 4-fold increase in total RLU generated, but no major change in fold-over-baseline and EC50 (**Ext. Fig.3**). The S153N mutant increases the dynamic range but does not have a major advantage in our system, and thus we opted to work with the non-mutated TEV219. We then moved on to adapt two other popular PPI assays, BRET and NanoBiT, for the NbV5 platform. Of note, a striking difference was discovered between BRET and NanoBiT at AT_1_R regarding the selective recruitment of β-arrestin isoforms. Interestingly, the preference for β-arrestin-1 over β-arrestin-2 is observed in the NanoBiT assay, while BRET^2^ and TANGO showed no disparity. The reason behind this observation is not clear. As shown in **Fig.6**, the size and orientation of the tag have a stronger effect on NanoBit than BRET^2^. This can be attributed to the fact that binary complementation is an interaction assay which cannot monitor spatial rearrangement. BRET^2^ is highly conformationally dependent, and the RET efficiency depends on the distance and spatial orientation to allow dipole-dipole coupling. One potential explanation is that the BRET^2^ signal measured is an average of the recruitment plus conformational changes (multi-states), while NanoBit only captures newly recruited β-arrestin or a single conformational state of the complex.

As aforementioned, the availability of the large-scale ORF library with a C-terminal V5-tag opens the possibility of performing mammalian functional proteomics. Its application will be extremely powerful to advance binary PPI mapping and is currently under investigation in our laboratory. This approach has the main advantages over other proteomics assay of not being limited to endogenous proteins, of being performed in living human cells, and of capturing ligand-dependent interactomes. For instance, a similar library has been recently used for the genome-wide profiling of human HOX proteins interactome using bimolecular fluorescence complementation (BiFc) assay. In said study, the authors employed a pooled library of 8200 ORFs and fused this set of ORFs to the C-terminal fragment of the blue fluorescent protein Cerulean, demonstrating this approach as a powerful tool to illuminate the binary PPI interactome. We strong believe that the NbV5:V5 system will make similar assays more accessible both economically and technically for the scientific community at large.

Finally, while the work presented herein utilized GPCRs to characterize the toolkit, we are confident that the NbV5:V5 system could be employed for probing any receptor-protein or protein-protein interaction^62^.

## CONCLUSION

Herein, we developed a new nanobody that recognizes the V5-tag epitope, which is a common peptide tag present in many expression vectors, including human genome-wide ORF library. Further, said nanobody was selected and maturated for its application in the intracellular environment in the form of an intrabody. The culmination of these efforts is our establishment of a versatile toolkit, named iBodyV5, for broad usage in NanoBiT, TANGO, BRET^2^ and microscopic imaging; even so, we are confident that our toolkit can further be expanded and other existing assays. In short, the iBodyV5 overcomes many limitations and allows the traceability of intracellular binding proteins with minimal disturbance.

## METHODS

### Cell culture

Human embryonic kidney 293T (HEK293T) and HT1080 cells were obtained from the American Type Culture Collection (ATCC) and maintained in Dulbecco’s modified Eagle’s medium (DMEM) supplemented with 5% fetal bovine serum (Fisher Scientific), 5% bovine calf serum (Fisher Scientific) and 1X Pen-Strep (100 U/ml penicillin, and 100 μg/ml streptomycin) (Fisher Scientific).

HEK293T cells stably expressing MOR-SmBiT (MOR-SmBiT/HEK293T) were generated by transduction with lentivirus particles, which had been produced in HEK293T cells by transiently co-transfecting the lentiviral packaging plasmid psPAX2 (Addgene #12260), a lentiviral vector encoding μ-OR-SmBiT (pLenti-Blast Addgene #17451), and the envelope plasmid pCMV-VSV-G (gift from Marceline Côté) at a 1:1:1 ratio using PEI transfection reagent. The following day, media was changed, and the supernatant was harvested at 48h post transfection. Cells were then transduced with lentivirus in standard growth media containing 5 μg/ml polybrene and the next day, selected with blasticidin at 5μg/m. All cells were cultured at 37°C in a humidified atmosphere containing 5% CO_2_.

### Plasmids and cloning

All plasmid DNA used in this publication were fully sequenced and are available upon request or through Addgene repository.

Plasmid encoding γ-Actin-V5 was extracted from the MISSION TRC3 Human LentiORF Puromycin library (MilliporeSigma).

V5-βArrestin1, V5-βArrestin2, βArrestin1-V5, βArrestin2-V5, βArrestin2-ALFA, ALFA-βArrestin2, βArrestin1-ALFA and ALFA-βArrestin1 were amplified by PCR, including the V5 tag within the primer, and subsequently cloned into pcDNA3.1^+^ at the HindIII-XbaI restriction sites.

GɑoA with internal V5-tag at position 92 was synthesized by IDT (Integrated DNA Technologies) and cloned at HindIII-XbaI sites in pcDNA3.1^+^.

BRET^2^ constructs: AT_1_R-RLuc8, μ-OR-RLuc8, NbV5-GFP2, GFP2-NbV5, βArrestin2-GFP2, βArrestin2-ALFA-GFP2 were amplified by PCR and cloned in pcDNA3.1^+^ using NEB HiFi DNA Assembly (New England Biolabs). Gβ3 and Gγ2-GFP2 were generously gifted by Dr. Asuka Inoue (TOHOKU University).

NanoBiT constructs: μ-OR-SmBiT and AT_1_R-SmBiT were amplified by PCR by including the SmBiT tag within the primer preceded by a (GGGS)_2x_ linker. NbV5-LgBiT, LgBiT-NbV5, NbALFA-LgBiT and LgBiT-NbALFA were amplified by PCR and cloned in pcDNA3.1^+^ using NEB HiFi DNA Assembly (New England Biolabs).

Gγ2-SmBiT was generously gifted by Dr. Asuka Inoue (TOHOKU University).

TANGO constructs: μ-OR-TANGO and AT_1_R-TANGO are from the original PRESTO-TANGO library ^29-31^. Aforementioned receptors fused to the TEV219 or TEV219 S153N mutant were cloned by PCR at restriction sites HindIII-BamHI in pcDNA3.1^+^-X-TEV219 vector. The S153N mutant was generated by site-directed mutagenesis.

Microscopy constructs: NbV5-eGFP was generated by PCR amplification of NbV5 and cloned into pEGFP-N1 (Clontech) at the HindIII-BamHI sites.

### Nanobody development

To identify nanobodies that bind to a linear target and that could be expressed from inside the cell (as intrabodies), phage display selection was conducted with the Nali-H1 synthetic library ^48^, by Hybrigenics Services (Paris, France), using a His-Halo protein fused with three successive V5-tags (Halo-3xV5). The naïve library was first depleted using another antigen fused in the same Halo vector and selected on magnetic streptavidin beads with the biotinylated Halo-3xV5. With the first round presenting a complexity of 3×10^6^ colonies, the DNA extracted from the first round were used to construct a Yeast-Two-Hybrid prey library by PCR and Gap repair. The VHH selected after one round of phage display against V5 tag-Biotin were cloned into the pP9 yeast prey vector, which is derived from the original pGADGH plasmid; the library had 2.6×10^5^ independent clones in yeast. A single V5 tag, as well as two tandem V5 tags, were cloned into pB27 as a C-terminal fusion to LexA (LexA-V5); pB27 is derived from the original pBTM116 plasmid ^63^. The constructs were verified by sequencing the insert and used as baits to screen the V5-specific VHH library For the screen, clones were vetted using a mating approach with YHGX13 (Y187 ade2-101::loxP-kanMX-loxP, matα) and L40ΔGal4 (mata) yeast strains as previously described ^50^. Moreover, the library has been screened at saturation by cell-to-cell mating ^50^. A total of 264 His+ colonies were selected on a medium lacking tryptophan, leucine and histidine supplemented with 0.5 mM 3-AT, obtaining 52 different VHH with redundancies from 1 to 37.

### Protein purification

The NbA1 was cloned into the expression vector pET26b (+) (Novagen) at NcoI-XhoI restriction sites to generate the pelB leader-NbA1-His_6_ construct. The plasmid was transformed in SHuffle T7 Competent E. coli (New England BioLabs), and the NbA1 was subsequently purified from the periplasm of SHuffle T7 cells. Bacteria were then grown at 30°C in Terrific Broth and, after reaching an OD600 of ∼0.6-0.8, were induced with 1mM IPTG (Isopropyl β-d-1-thiogalactopyranoside, ThermoFisher) at 25°C for approximately 16 hours. Bacteria were then pelleted and resuspended in a solution of lysis buffer (0.5 M sucrose, 0.2 M Tris pH 8, 0.5 mM EDTA) and water at a ratio of 1:2 to create an osmotic shock. The lysate was then frozen and thawed for more efficient purification. Followingly, the mixture was stirred for 45 minutes at 4 °C and brought to a concentration of 150 mM NaCl, 2 mM MgCl_2_, and 20 mM imidazole and centrifuged at 20,000xg for 30 min at 4 °C. The supernatant was then filtered through a 0,22 μm filter. After filtration, the supernatant was added to a gravity column containing 4 mL of Ni-NTA (Qiagen). Beads were washed with a high salt buffer (20 mM HEPES pH 7.5, 500 mM NaCl, 20 mM Imidazole) and washed three times with low salt buffer (20 mM HEPES pH7.5, 100 mM NaCl, 20 mM imidazole). The nanobodies were then eluted (20 mM HEPES pH 7.5, 100 mM NaCl, 400 mM imidazole) and dialyzed into physiological buffer solution (10 mM HEPES pH 7.4, 140 mM KCl, 10 mM NaCl) and purified using FPLC (Fast protein liquid chromatography, AKTA GE) on an S75 prep size-exclusion column.

### Crystallization, data collection and structure determination

Purified NbA1 (30mg/mL) was incubated with the V5 peptide in a 1:3 ratio (protein: peptide). The protein complex was crystallized via the sitting drop vapour diffusion method at 4°C with a mother liquor composed of Bis-Tris pH 6.5, 19% (w/v) PEG 3350 and 20% (v/v) ethylene glycol. The crystals were flash-frozen in liquid nitrogen, and a full data set was collected using a Rigaku MicroMax-007HF equipped with a copper anode. Images were collected using an R-Axis IV++ detector (Rigaku) and processed using Structure Studio (Rigaku). The structure was solved by molecular replacement using the structure of NbALFA (PDB 6I2G) as search model and Phaser ^42^. Following several rounds of NbA1 building and refinement using COOT and Phenix, respectively, V5 was built in the positive Fourier map. The model was completed by adding the molecules and truncating side chains for which no electronic density could be observed. Ramachandran statistics: Non-glycine Ramachandran outliers; 0%, Non-glycine Ramachandran favored; 100%,Molprobity score : 1.65. Statistics of data collection and refinement are summarized in Table1. All structural figures were prepared in PyMOL.

### Isothermal titration calorimetry (ITC)

ITC experiments were performed on a VP-ITC calorimeter (MicroCal, Northampton, MA) by injecting the V5-peptide (521 μM) into the cell containing the NbV5 (38.7 μM) in HBS buffer solution (10 mM HEPES pH 7.4, 140 mM KCl, 10 mM NaCl) with or without 5 mM DTT. The assay was performed at 19°C and analyzed using the Origin software (OriginLab Corporation, Northampton, MA).

### Nanobody maturation

To optimize the NbA1 sequence and potentially increase the affinity of the antibody towards V5, Rosetta single-state design protocol was performed using the Rosetta Software Suite on the crystal structure of the NbA1 complex. The structure of NbA1 was prepared for antibody affinity maturation for Rosetta by manually editing the PDB file in PyMOL. PyMOL command prompts were used to delete the unwanted water molecules and all non-essential ligands and chains. An extra processing step was also performed to remove any protein atoms that are not involved in the antibody-antigen interface; chains were also renamed and reordered to help differentiate the antibody residues from the antigen residues. Next, a resfile (python script) was generated to identify the residues that were within a distance of specified residues that define the NbA1 protein interface. The side-chain conformations were optimized using the repacking and relaxing feature in Rosetta protein design, which was performed to minimize backbone phi-phi angles to relieve small clashes between sidechains. The relaxed model was then used to generate ten designed models through RosettaScripts XML file, which contains a design protocol that uses a single round of fixed backbone design. As a control, the same protocol was repeated to generate ten control models without designing any residues. These control models were generated to compare the scores and the binding energies of the designed models to the native sequence during the analysis stage. The designed sequences were then analyzed by looking at the score, binding energy, and the binding density of the models. The analysis of the metrics was plotted using RosettaScripts which plots the score and the binding energy of the deigned models against the control models. Subsequently, specific mutations were identified that resulted in the improvement of the NbA1 complex based on corresponding findings from functional assay experiments (as shown in Figure 1). Finally, a sequence logo was generated from the designed models in order to determine which mutations were made and their frequencies.

### BRET^2^ assay

HEK293T cells were plated in 6-well plates at 1.2×10^6^ cells and subsequently transfected with BRET^2^ constructs (at ratios indicated in the Figure legends) at a total of 3 μg of DNA per well. Transfected cells were detached and seeded on PLL-coated white 96-well assay plates (ThermoFisher). The following day, spent medium was removed and replaced with 60 μL of 1X HBSS buffer, followed by the addition of 10 μL of Coelenterazine 400a (Nanolight Technologies) at 50 μM to each well, for a final concentration of 5 μM. After incubating the plates away from light for 8 minutes, 30 μL of serial dilutions of agonists at 3X concentration was added. Plates were subsequently read 4 times after 2 minutes, 10 minutes, 20 minutes, and 30 minutes using the Hidex Sense Beta Plus microplate reader (Gamble Technologies) with 405 nm (RLuc8-Coelenterazine 400a) and 500 nm (GFP2) emission filters, at 1 second/well integration times. Figures shown in the manuscript account for the reads after 20-minute incubations with agonist (and correspondingly, approximately 30-minute incubations with Coelenterazine 400a). Data were extracted using the integrated software and subjected to non-linear least-squares regression analysis using the sigmoidal dose-response function provided in GraphPad Prism 9.0. Data of 3 or 4 independent experiments (N=3 or 4) performed in quadruplicate are presented as BRET^2^ ratio (acceptor/donor) as indicated in figure legends.

### NanoBiT assay

μ-OR-SmBiT/HEK293T and HEK293T cells were plated in 6-well plates at 1.0×10^6^ cells and then transfected with the NanoBiT constructs the next day (ratio and constructs in Figure legends) at a total of 3 μg of DNA per well. Transfected cells were detached and seeded on PLL-coated white 384-well assay plates (ThermoFisher) in starvation media (DMEM, 1% FBS, 1x Pen-Strep), The next day, media was removed and replaced with 20μL of 1X HBSS buffer containing 5 μM furimazine and incubated for a total of 10 minutes at room temperature before reading on Fluorescent Imaging Plate Reader (FLIPR) Tetra system (Molecular Devices). Baseline measurements were initially read before drugs were added into their respective wells (concentration and different drugs in Figure legends). The subsequent changes in relative luminescence signals (RLU) were recorded over time. Data were extracted using the integrated ScreenWorks software and subjected to non-linear least-squares regression analysis using the sigmoidal dose-response function provided in GraphPad Prism 9.0. Data of 3 or 4 independent experiments (N=3 or 4) performed in quadruplicate are presented as Relative Luminescence Units (RLU) or normalized as indicated in figure legends.

### TANGO assay

HTTL (HEK293T stably expressing a luciferase reporter gene under the TRE-Tight promoter) cells, an in-house developed reporter cell line (to be published), were seeded in 6-well plates at 1.2×10^6^ cells and were transfected with TANGO-ized constructs using the PEI precipitation method. Twenty hours later, the transfected cells were plated in in DMEM supplemented with 1% dialyzed FBS into PLL coated 384-well white clear bottom cell culture plates at a density of 20,000 cells/well in a total volume of 40 μL for 5 hours to ensure proper attachment of cells. Agonist solutions, previously prepared at 3X concentration in sterilized assay buffer (20 mM HEPES, 1× Hanks’ balanced salt solution (HBSS), pH 7.40), were added to the cells at 20 μL per well. Following overnight incubation, media was removed and 20 μL per well of homemade luciferase detection reagent (108 mM Tris–HCl; 42 mM Tris-Base, 75 mM NaCl, 3 mM MgCl_2_, 5 mM Dithiothreitol (DTT), 0.2 mM Coenzyme A, 0.14 mg/ml D-Luciferin, 1.1 mM ATP, 0.25% v/v Triton X-100, 2 mM Sodium hydrosulfite) was added to all wells. After 10 minutes of incubation in the dark at room temperature, plates were read using the Hidex Sense Beta Plus microplate reader (Gamble Technologies). Data were subjected to non-linear least-squares regression analysis using the sigmoidal dose-response function provided in GraphPad Prism 9.0. Data of 3 or 4 independent experiments (n=3 or 4) performed in quadruplicate are presented as Relative Luminescence Units (RLU) or normalized as indicated in figure legends.

### Immunofluorescence

HT1080 were seeded in a chambered coverslip with 8 individual wells and high walls in complete medium to obtain a 50% confluency the following day. The next day, cells were transiently cotransfected using JetPRIME (Polyplus Transfection) with γ-actin-V5/pLX307 or pcDNA3 and NbV5-eGFP/pcDNA3.1+. Twenty-four hours post-transfection, cells were fixed with 4% paraformaldehyde, washed and then counterstained with Hoechst 33342 stain solution. Cells were imaged on a Zeiss Axio Observer D1 fluorescence microscope at 100X, and the images were analyzed using ImageJ software.

## FIGURE LEGENDS

**Extended Fig.1:**
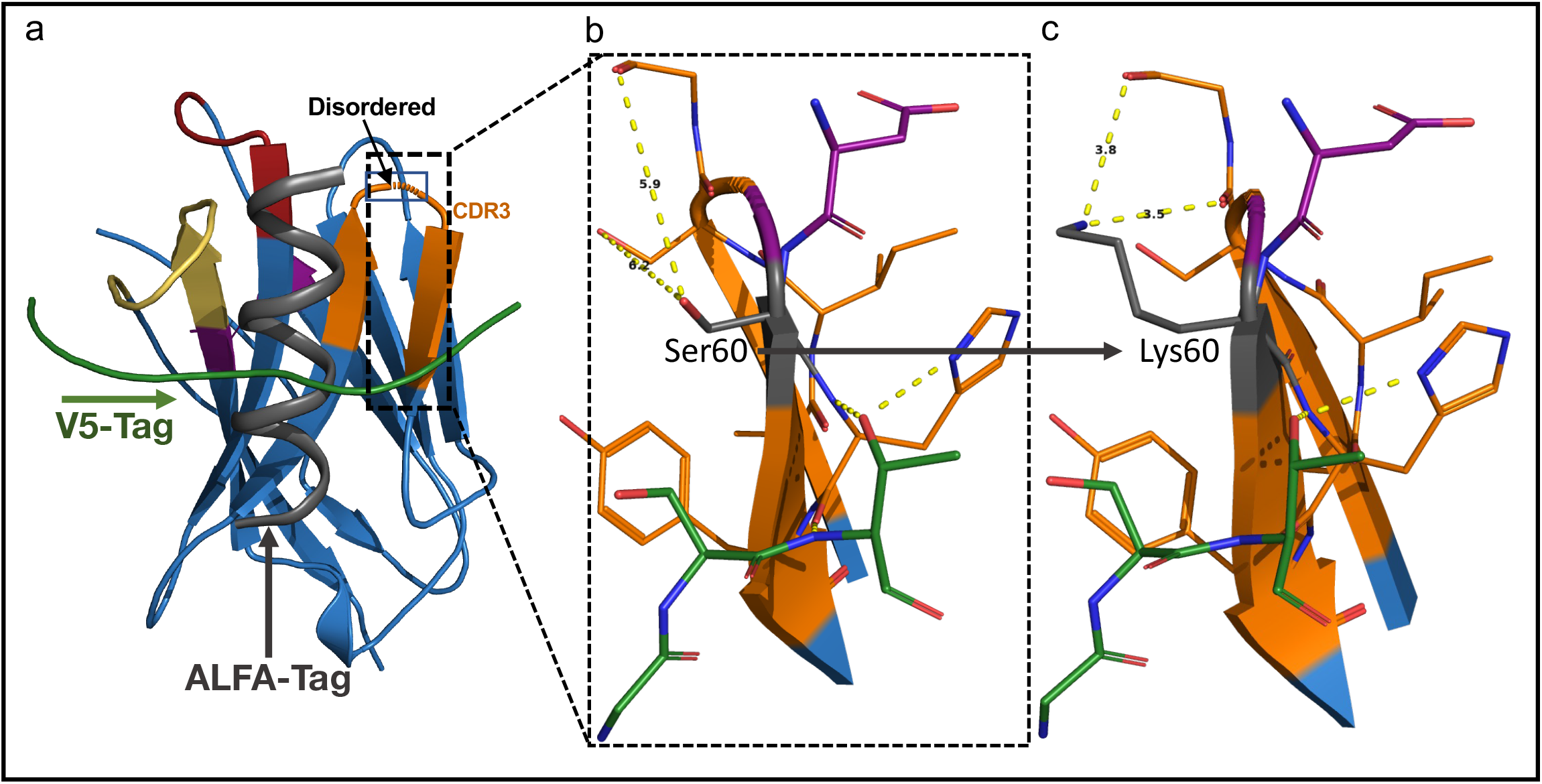
**a**, Overlay of the NbA1:V5 structure with the NbALFA:ALFA (PDB: 6I2G) structure. The NbALFA structure was omitted for simplicity and to present the binding pose of the ALFA-tag, which is at 90 degrees compared with the V5-tag. Using *in silico* maturation and functional studies assessment, the mutations ΔAsp^59^, Ser^60^Lys were discovered to substantially improve the behavior of the nanobody. **b**, Close-up view of the Asp^59^ and Ser^60^ of the CDR2 loop that are juxtaposed to the disordered tripeptide RQG. The Ser^60^ is not involved in any interaction and the distance with the closest residues is shown with a dash line. **c**, It is hypothesized that mutations ΔAsp59 and Ser^60^Lys could favorize new intramolecular polar interactions within the CDR2 loop which might stabilize it by reducing entropy.

**Extended Fig.2:**
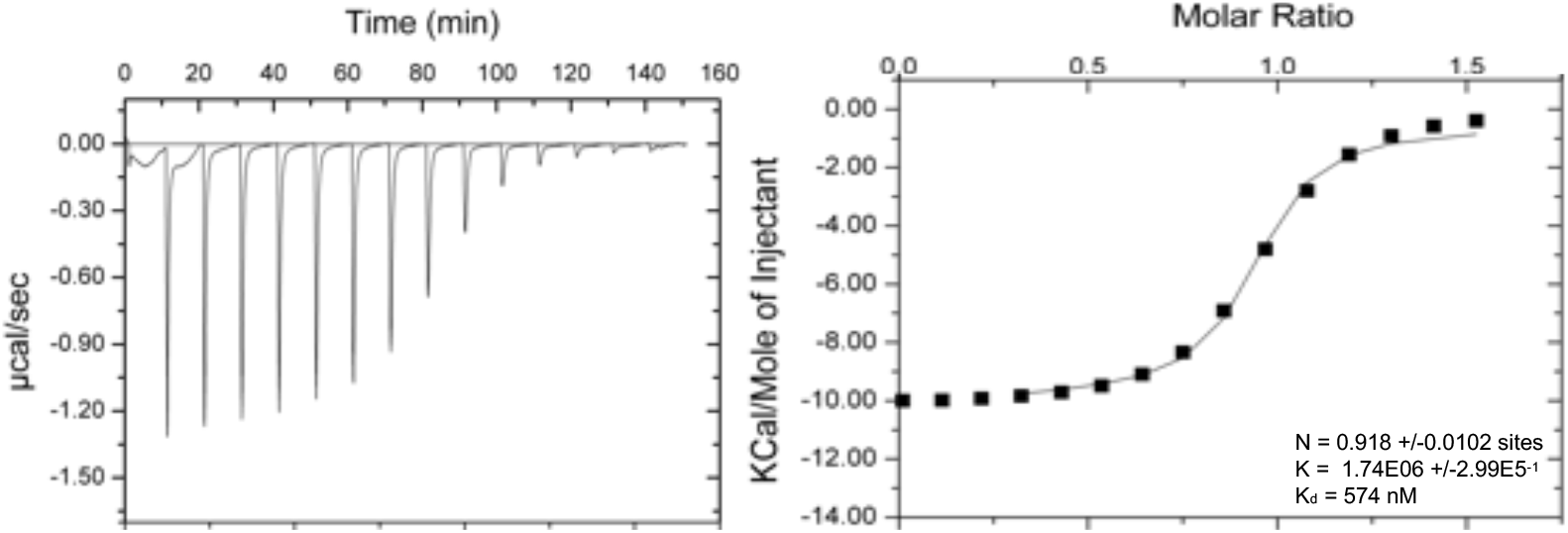
Isothermal titration calorimetry curve of V5-peptide tag titrated into NbV5 in the presence of DTT and corresponding ITC values. Titrations were performed in duplicate. S.D. represents the standard deviation between the two experiments.

**Extended Fig.3:**
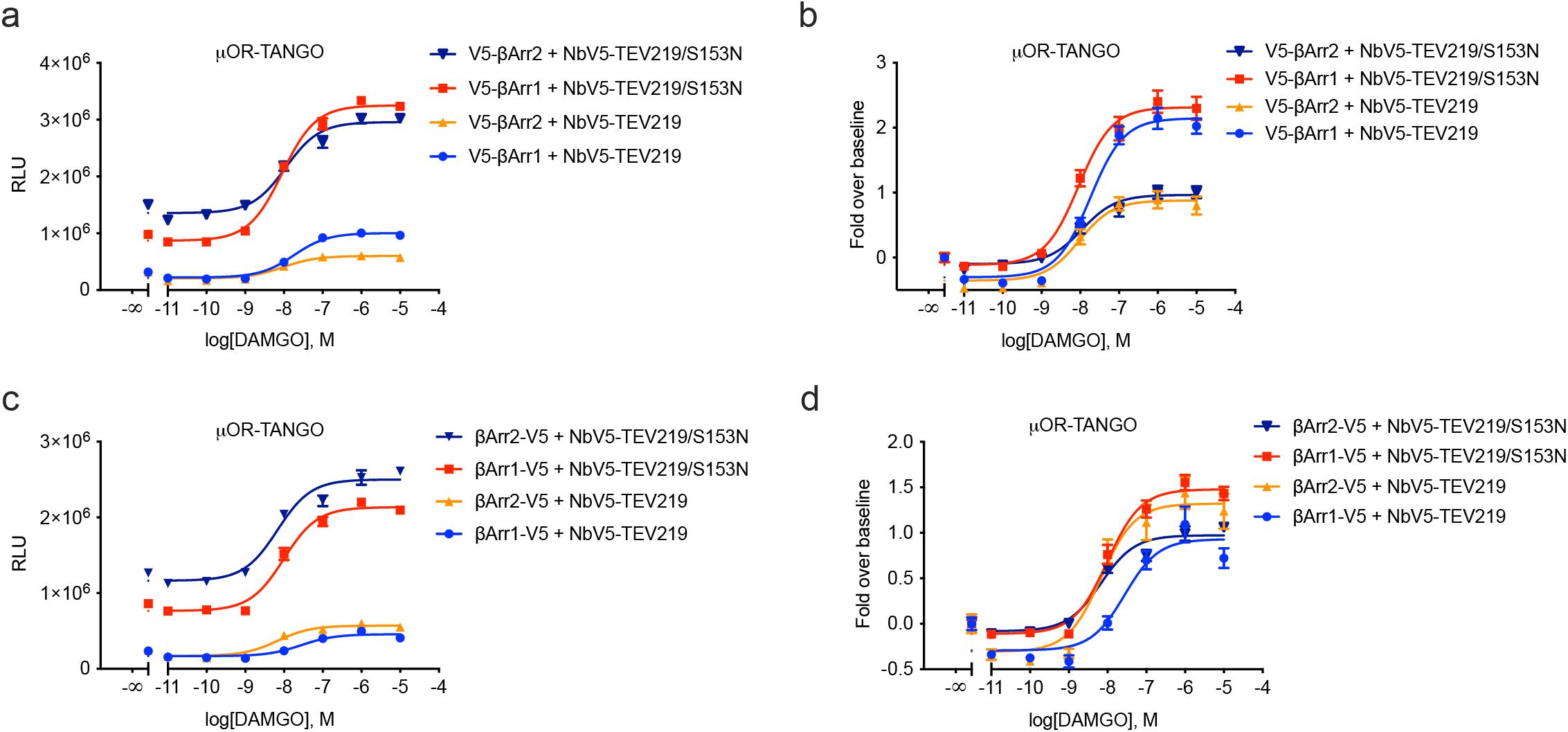
The new S153N variant of the TEV219 was tested to assess whether the catalytic activity (k_cat_) is a limiting factor in our system. As shown in (**a-d**) for the μ-OR, the S153N mutant exhibits a strong increase in total RLU, which correlates with an increased catalytic rate, but the fold-over baseline as well as the EC50 are not improved. Dose-response curves were built using XY analysis for non-linear regression curve and the 3-parameters dose-response stimulation function. All error bars represent SD (n = 3 technical replicates).

## ACKNOWLEDGMENTS

KM is supported by a graduate scholarship from the Natural Sciences and Engineering Research Council of Canada. MZ is supported by the Alexander Graham Bell Canada Graduate Scholarships-Doctoral Program (CGS-D3) from Natural Sciences and Engineering Research Council of Canada. This work was supported by the Canadian Institutes of Health Research (CIHR grant #MOP142219) and Natural Sciences and Engineering Research Council of Canada (NSERC RGPIN-2017-06151). We would like to acknowledge technical support from the Protein Biophysics Core Facility and uOttawa Cell Biology and Image Acquisition Core Facility. We would like to thank Dr. Asuka Inoue for his generous gift of different plasmids as highlighted in the Material and Methods. Thank you to Hybrigenics (France) for performing the nanobody selection and Y2H.

## AUTHOR CONTRIBUTIONS

PMG conceived the overall concept; MZ, KM, AV and GL performed all functional experiments and analyzed the data. MZ and PMG wrote the manuscript. SP performed the *in silico* maturation of the nanobody. JFC, SS, MJ provided the technical support for the protein crystallization and data refinement and ITC.

